# Recomb-Mix Cloud: a cost-efficient and scalable cloud-friendly pipeline for local ancestry inference

**DOI:** 10.64898/2026.09.28.755192

**Authors:** Yuan Wei, Degui Zhi, Shaojie Zhang

## Abstract

Local ancestry inference (LAI) reveals population compositions of an individual’s chromosomes, which empowers population genetics and genetic association studies of diverse populations. The recent availability of large genomic datasets from biobanks offers unprecedented opportunities for LAI. However, such datasets are typically only accessible from cloud-based platforms, and the lack of efficient and cost-effective tooling makes LAI prohibitive for resource-constrained researchers. Here, we develop Recomb-Mix Cloud, a cloud-ready pipeline for LAI based on Recomb-Mix, a recently introduced efficient LAI method. Recomb-Mix focuses on individual variants while maintaining a lower computational burden than other LAI methods focusing on genomic windows. Recomb-Mix Cloud offers a cloud-friendly complete pipeline from panel sample selection, marker alignment, compact reference panel construction, Recomb-Mix threading, and generating LAI calls in common data formats. Recomb-Mix Cloud also offers a set of pre-computed compact population reference panels constructed based on data from the Human Genome Diversity Project and the 1000 Genomes Project. Our results showed Recomb-Mix is about one and two orders of magnitude cheaper and faster than FLARE and RFMix, two popular LAI methods, while offering largely consistent LAI calls on the All of Us Researcher Workbench. The estimated overall compute cost per All of Us sample for whole-genome LAI analysis using Recomb-Mix Cloud with a pre-computed reference panel is just $0.01, compared with $0.43 for FLARE and $4.58 for RFMix. Therefore, Recomb-Mix Cloud is well-positioned to democratize local ancestry-based genetics research on modern biobank-sized data. The Recomb-Mix Cloud pipeline and the compact reference panels are available at https://github.com/ucfcbb/Recomb-Mix.

## 1 Introduction

Many biobank datasets are now available through cloud services, such as the All of Us Researcher Workbench (AoURW) [4], and the UK Biobank research analysis platform [15]. These datasets involve a diverse group of millions of individuals and aim to advance research in human genetics, including personalized health and precision genomic medicine. Individuals of admixed ancestry have received increased attention with the rise of ancestry-aware analyses, which provide deeper insights into sub-population structure and health outcomes. Many methods leveraging local ancestry information have been developed for ancestry-aware genomic analysis. Tractor delivered highly accurate genetic association signals, enhancing the identification of gene variants linked to increased disease risk in admixed populations [1]. Tractor also produced ancestry-specific allele frequencies from the Genome Aggregation Database (gnomAD) [10], enhancing the interpretation of genetic variants and the genetic architecture of complex traits across diverse genetic ancestry groups [18]. Tractor-Mix enabled local ancestry-informed genome-wide association studies (GWAS) in admixed cohorts, effectively accounting for relatedness among individuals [29]. Multiple approaches to calculating polygenic risk scores (PRS) have incorporated local ancestry information, resulting in improved predictive accuracy for certain polygenic traits in admixed populations [23, 28, 31, 35]. A transcriptome-wide association study pipeline for admixed cohorts was developed to leverage local ancestry information alongside summary-level expression quantitative trait locus data. This approach achieved superior performance and identified unique gene-trait associations not detected by traditional local ancestry-unaware approaches [16]. An ancestry-specific expected genetic relationship matrix framework was proposed to reveal ancestry-specific structures in admixed populations. By integrating local ancestry information into an ancestral recombination graph-based model, this approach improved the resolution in uncovering ancestry-specific fine structure through principal component analysis and uniform manifold approximation and projection [30]. A positive selection analysis integrating local ancestry information was developed to determine whether signals of adaptation resulted from pre- or post-admixture selection for populations with complex demographic histories [9]. Admixture mapping-based association studies leveraging local ancestry information were performed, enabling the characterization of cross-population phenotypic differences [21] and the discovery of complex trait associations [13] within the All of Us Research Program.

LAI methods have been continuously developed since the early 2000s, with new approaches still emerging [27]. The foundation of LAI lies in the Li and Stephens (LS) Hidden Markov Model (HMM) [20], which conceptualizes alleles or segments as observed variables and their ancestral origins as unobserved variables. LAI methods are generally categorized into window-based or site-based methods. Window-based approaches leverage linkage disequilibrium information to infer ancestry. Recent and widely used examples include RFMix [22], G-Nomix [17], and Orchestra [19]. In contrast, site-based methods focus on locus-specific ancestry signals, as seen in methods such as FLARE [7], SparsePainter [33], and Recomb-Mix [32]. Recomb-Mix was designed for whole-genome sequencing data and for cases of recent inter- and intra-continental admixture events. Additionally, it accommodates compact reference panels that are compressed at the haplotype level, further improving efficiency.

While recent population genetic studies have increasingly incorporated local ancestry inference (LAI) for admixed individuals, several challenges persist, particularly with data hosted on cloud platforms. One major issue is the high cost associated with running analyses in these environments. For example, a previous study using the conventional RFMix method to perform local ancestry calls on the All of Us (AoU) data for PRS calculations highlighted these expenses: for a dataset of 53,944 individuals and 629,877 variants, local ancestry inference by RFMix incurred the highest cost ($400), compared to $60 for variant selection and $120 for phasing [35]. Note that the AoU version 8 phased genomic dataset contains 658,219,451 variants. Running LAI on this dataset using RFMix may incur significant cost. Additionally, reference panels for LAI are often not readily accessible, requiring users to prepare their own panels to suit their study objectives. While some LAI tools provide reference panels or pre-trained models, these resources are typically tailored to a specific context and may not address the diverse requirements of all users. For example, target individuals and available reference panels are in different reference builds, and training a custom model is compute-intensive and often infeasible in the cloud due to platform memory constraints [13]. Furthermore, many LAI tools generate biased results, often overestimating or underestimating specific ancestries because of substantial discrepancies in sample proportions within the reference panel. For example, certain non-European local ancestry labels were excluded from the results due to the small sample sizes in the reference panel used for the ancestry inference [13]. Last but not least, there is no widely accepted community standard for LAI output formats. Each LAI tool has its own output format, which complicates the studies since it requires users to develop custom scripts to convert outputs for their downstream analyses, making the overall process time-consuming.

Here, we introduce a cloud-based local ancestry inference solution that offers a user-friendly, Recomb-Mix-centralized LAI pipeline alongside an extensive collection of compact reference panels. Our proposed pipeline significantly reduces both computational costs and cloud storage demands compared to conventional pipelines when conducting high-accuracy local ancestry calls on cloud-hosted datasets. By providing a diverse set of pre-computed reference panels in a compact format, our solution accommodates a wide range of analysis needs. Ultimately, we aim to lower barriers for researchers accessing LAI information from cloud data and to promote its integration into population genetic studies of admixed populations.

## 2 Materials and methods

### 2.1 Recomb-Mix Cloud

The Recomb-Mix Cloud pipeline is a comprehensive, cloud-friendly pipeline that guides users through the steps of local ancestry inference from the raw target dataset. The pipeline encompasses sample selection, marker alignment, construction and utilization of compact reference panels, local ancestry inference powered by Recomb-Mix, and generation of LAI calls in space-efficient and common data formats. The pipeline simplifies the LAI process for users, reduces computational cost, and accelerates user studies by ensuring compatibility with the expected input data format for downstream analysis.

Recomb-Mix Cloud streamlines the LAI process for individual samples in a cloud platform. The pipeline relies on Recomb-Mix [32], an efficient and accurate LAI method designed for whole-genome sequencing data, to make local ancestry calls. Recomb-Mix is a site-based LAI method that approximates the LS HMM. By leveraging discrete ancestry-informative markers, Recomb-Mix effectively captures strong ancestry signals, enhancing the method’s accuracy and power. The LS HMM approximation uses a haplotype-level compressed reference panel to simplify path penalty calculations, resulting in substantial efficiency gains. Thus, Recomb-Mix is well-suited as an LAI method for obtaining local ancestry calls from data in cloud computing environments.

The pipeline accepts genotype files for admixed individuals and the reference panel, ancestry labels for the reference panel, and a recombination map file as input. It first filters sites and samples using PLINK 2 [24], retaining only the targeted admixed individuals and the shared biallelic sites present in both the admixed and reference panels. While alternative software options are available here, PLINK 2 is chosen because it supports a wide range of popular genomic data input formats, including BED, PED, or PGEN (PLINK binary or text genotype), GEN or BGEN (Oxford-format binary or text genotype), and VCF (variant call). Next, Recomb-Mix is invoked to generate local ancestry calls across the chromosome. The resulting output file provides local ancestry intervals for each admixed individual’s haplotype using run-length encoding. While the default run-length encoding format efficiently stores LAI calls, a conversion utility enables users to convert the output file into ancestry dosage or ancestry state formats as needed for downstream analysis. Overall, the proposed pipeline significantly reduces both computational costs and cloud storage demands compared to conventional pipelines when conducting high-accuracy local ancestry calls on cloud-hosted datasets. Figure 1 illustrates the advantages of Recomb-Mix Cloud compared to a conventional pipeline for LAI with AoU genomic data. Recomb-Mix Cloud achieved significantly lower computational costs and reduced cloud storage requirements, while delivering superior accuracy in local ancestry inference. Note that the AoURW is just an example, and the Recomb-Mix Cloud pipeline is compatible with various cloud environments. A step-by-step tutorial for running the Recomb-Mix Cloud pipeline, including an example script for downloading software, filtering markers and samples, and performing local ancestry inference, is available on GitHub here: https://github.com/ucfcbb/Recomb-Mix/blob/main/Pipeline.md.

**Figure 1:**
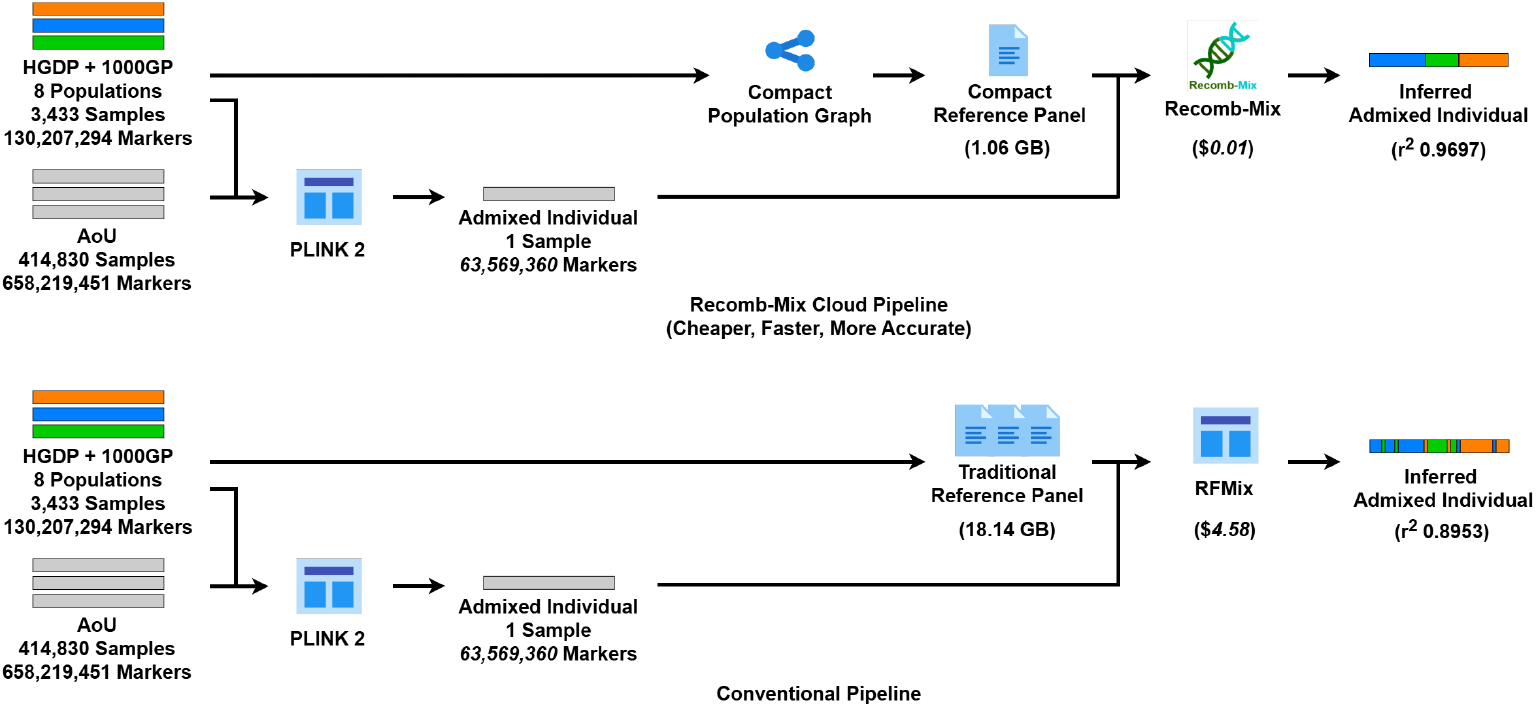
Recomb-Mix Cloud pipeline substantially reduces computational costs and cloud storage requirements while achieving superior local ancestry inference accuracy for the All of Us (AoU) genomic data, compared to the resource-intensive conventional pipeline. The sizes of reference panels were ones containing eight ancestries and constructed from the Human Genome Diversity Project (HGDP) and the 1000 Genomes Project (1000GP), phased and imputed by Beagle. The number of markers and overall costs of LAI with Recomb-Mix and RFMix (in *Italics*) were estimated on the AoU data for 200 individual samples. The mean diploid squared Pearson correlation coefficient *r*^2^ values were calculated based on a simulated 3-way inter-continental 15-generation 500-reference Chromosome 18 dataset, averaged by 200 admixed individuals.

### 2.2 Reference panels

A robust reference panel is essential for achieving accurate LAI results. The Recomb-Mix Cloud pipeline provides reference panels built from the Human Genome Diversity Project (HGDP) [3] and the 1000 Genomes Project (1000GP) [8] in the GRCh38 human genome reference build, ensuring comprehensive representation of ancestries. Creating a compact reference panel involves two key steps: harmonizing the source data and constructing the compact reference panel (Figure 2). To create the compact reference panels, the HGDP and 1000GP datasets were merged, phased, and imputed using Beagle 5.4 [5, 6], resulting in 130,207,294 sites. Ancestry labels for each sample were determined based on the population and sub-population labels from their respective original datasets. The reference panels were then transformed into compact population graphs by collapsing sites with distinct allele values and ancestral population labels, and stored as Variant Call Format (VCF) [12] files. We provide several compact reference panels available for the populations frequently represented in downstream analyses: two-way AFR-AMR, two-way AFR-EUR, three-way AFR-EAS-EUR, three-way FIN-GBR-TSI, seven-way AFR-AMR-EAS-EUR-OCE-SAS-WAS, and eight-way AFR-AMR-CAS-EAS-EUR-OCE-SAS-WAS. The reference panels are constructed using the GRCh38 [25] human genome reference. Additionally, equivalent panels for the GRCh37 [11] build are provided to ensure compatibility with the substantial number of legacy systems and datasets that still rely on the GRCh37 build. The genome coordinates were lifted over from the GRCh38 build using CrossMap [34]. A utility to convert original reference panels into compact ones is also available to the users, enabling reference panel customization based on users’ specific study needs.

**Figure 2:**
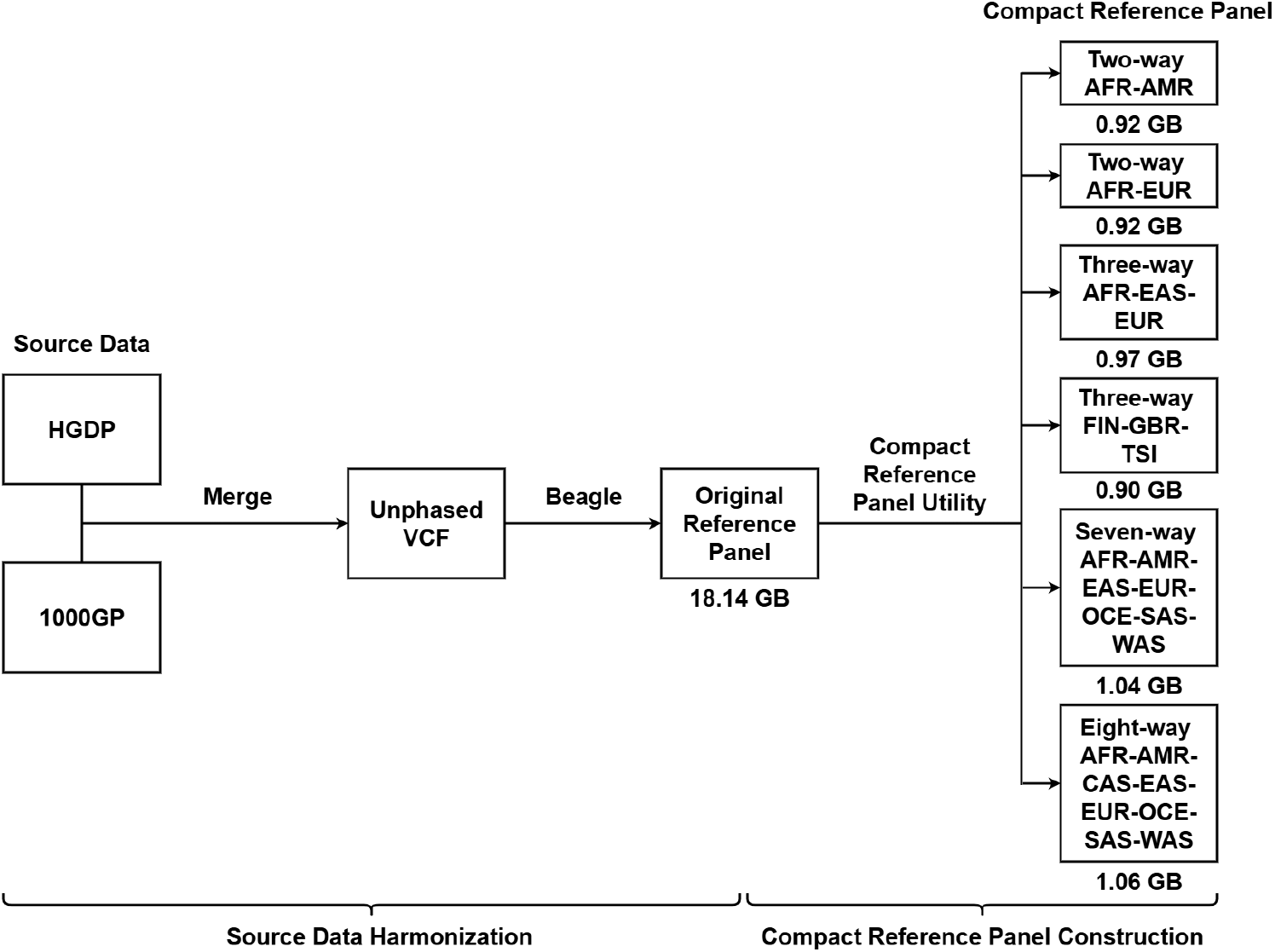
Overview of creating compact reference panels. The process consists of two main steps: source data harmonization and compact reference panel construction. The source data are the Human Genome Diversity Project (HGDP) and the 1000 Genomes Project (1000GP). The available populations in the compact reference panels are African (AFR), Admixed American (AMR), Central Asian (CAS), Eastern Asian (EAS), European (EUR), Finnish in Finland (FIN), British in England and Scotland (GBR), Oceanian (OCE), South Asian (SAS), Toscani in Italia (TSI), and Western Asian (WAS).

### 2.3 LAI output formats

By default, Recomb-Mix outputs local ancestry calls using run-length encoding, which significantly reduces storage requirements compared to other LAI output formats. RFMix produces its ancestry calls as a matrix with dimensions equal to the number of individual haplotypes by the number of windows. The window is an input parameter indicating a genomic interval that is the smallest unit of an ancestry tract during the inference. FLARE generates a new VCF file with ancestry labels inserted alongside genotype values. RFMix and FLARE output formats require 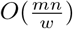 and *O*(*mn*) storage space, where *m* is the number of individual haplotypes, *n* is the number of sites, and *w* is the window size. In contrast, run-length encoding is much more space-efficient for local ancestry calls. This format includes three sections: Ancestry, Position, and Result. The Ancestry section contains the population labels and their IDs, with one population per line, in tab-delimited format. The Position section lists physical positions in ascending order, with one position per line. The Result section contains the inferred ancestry intervals for each query individual haplotype, in tab-delimited format. Each line represents the result of a single individual haplotype, starting with its ID, followed by a list of intervals, each sorted in ascending order, with three fields: start physical position, end physical position, and inferred ancestry ID. The inferred ancestry ID is a zero-based index of the population label, and the corresponding population label can be found in the Ancestry section by its population ID. The schema definition is in Table 1. The required space of run-length encoding format is substantially small, at *O*(*p* + *n* + *m*), where *p* is the number of ancestries.

**Table 1:** The run-length encoding format schema.

| Section | Element | Type | Description |
| --- | --- | --- | --- |
| Ancestry | population_label | string | Name of a population |
| Ancestry | population_id | integer | Unique identifier of a population |
| Position | physical_position | integer | Base coordinate of a chromosome |
| Result | individual_haplotype_id | string | Identifier of an individual haplotype |
| Result | start_physical_position | integer | Inclusive start physical position of an interval on a chromosome |
| Result | end_physical_position | integer | Inclusive end physical position of an interval on a chromosome |
| Result | ancestry_id | integer | Inferred population_id for an interval on a chromosome |

While run-length encoding format is space-efficient, many downstream analyses, including local ancestry-aware association studies, prefer formats that allow direct access to local ancestry information at the site level. For instance, Tractor uses an ancestry dosage-like format, in which each row represents a site and each column indicates the count of a given ancestry for an individual at that site. In this format, each ancestry’s count is 0, 1, or 2, and the total ancestry count per site for each diploid individual is always 2 (reflecting two haplotypes per individual). The storage requirement for this format is *O*(*pmn*). To address the need for more accessible formats, a utility is provided to convert run-length encoding format to other commonly used formats, such as ancestry dosage and ancestry state (Figure 3). See Supplementary Tables S1 and S2 for the schema definitions. This flexibility ensures that users can easily integrate our local ancestry calls from the Recomb-Mix Cloud into their downstream analysis pipelines.

**Figure 3:**
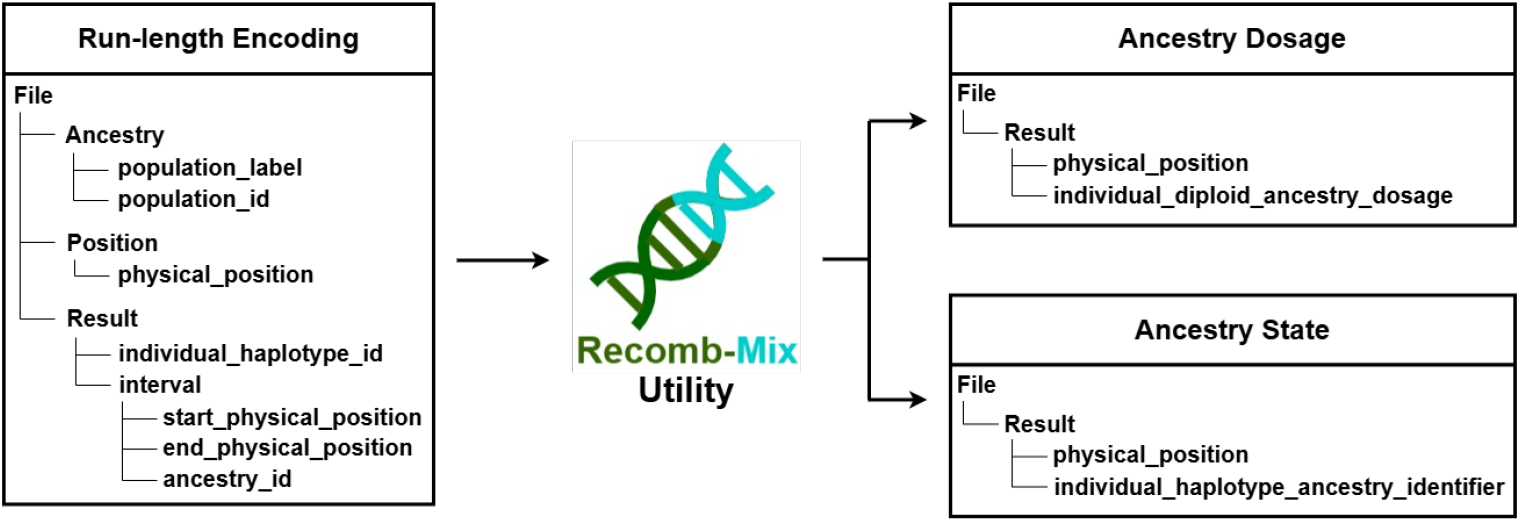
LAI output format conversion. The Recomb-Mix Utility converts the run-length encoding format to the ancestry dosage or the ancestry state format.

## 3 Results

### 3.1 Accuracy

We evaluated the LAI performance of Recomb-Mix, FLARE [7], and RFMix [22] through simulations. We used the same simulation pipeline (described in [32]) to generate 200 admixed individuals from three-way inter-continental (African, East Asian, European) Chromosome 18 data. The simulations included varying reference panel sizes (100, 250, 500, and 1,000) and different numbers of generations since the admixture event (15, 50, 100, and 200). For accuracy assessment, all sites from the simulated datasets were included, and the mean diploid squared Pearson correlation coefficient *r*^2^ across all ancestries was used, which is the same metric used in the FLARE paper: “We calculated the squared Pearson correlation of the estimated and true diploid ancestry dose across all individuals and sites. Separate *r*^2^ values were calculated for each ancestry and overall reported values are the unweighted mean *r*^2^ across all ancestries” [7]. Notably, this definition of *r*^2^ differs from that used in LAMP-LD [2], which was previously reported for benchmarking LAI methods [32].

In all seven simulated scenarios involving three-way intercontinental Chromosome 18 datasets, Recomb-Mix consistently outperformed FLARE and RFMix. Specifically, it had superior results on a 15-generation dataset with reference panel sizes of 100, 250, 500, and 1,000, as well as on a dataset with 500 reference samples and generations of 50, 100, and 200 (see Supplementary Table S3). These results highlight Recomb-Mix’s robust performance in challenging LAI scenarios. For example, it surpassed other LAI methods when dealing with a small reference panel (e.g., the REF 100 case) and a high number of generations since admixture (e.g., the GEN 200 case). This makes Recomb-Mix well-suited not only for common LAI analyses but also for studies involving minor populations with small reference-panel sample sizes and distant admixture events.

### 3.3 Disk usage

We compared the sizes of the eight-way inter-continental original and compact reference panels, and the output files for the case of running LAI on a simulated three-way inter-continental 15-generation 100-reference Chromosome 18 dataset for 200 admixed individuals with Recomb-Mix, FLARE, and RFMix. The overall size of the compact reference panels for the whole genome is only 1.06 gigabytes, which is 5.84% of the size of the original reference panels (18.14 gigabytes). See Supplementary Table S4 for the summary of statistics for the reference panels. As for the LAI output formats, we found that the run-length encoding output has the smallest file size (77 kilobytes, see Table 2) on the simulated three-way inter-continental 15-generation 100-reference Chromosome 18 dataset, while preserving 100% of the original result (i.e., local ancestry calls at each physical position). FLARE’s result also preserves all the ancestry calls; however, the way it stores as a VCF file costs a lot of disk space even though the file is compressed. RFmix’s matrix file size is small, but it does not retain all physical position information and requires the original query file to recover the ancestry calls for each physical position. Ancestry dosage focuses on the diploid ancestry counts, so it discards the ancestry information on the haplotype level. Ancestry state contains all ancestry calls for all positions in a matrix format, which is convenient for downstream analysis; however, it requires a large amount of disk space.

**Table 2:** The data retention, compression, and file sizes (kilobytes) of the 200-individual LAI output formats on the simulated three-way inter-continental 15-generation 100-reference Chromosome 18 dataset.

| LAI Output Format | Data Retention | Compression | File Size |
| --- | --- | --- | --- |
| Recomb-Mix (Run-length encoding) | Lossless | No | 77 |
| FLARE (VCF) | Lossless | Yes | 13,215 |
| RFMix (Matrix) | Lossy | No | 242 |
| Ancestry Dosage | Lossy | Yes | 3,010 |
| Ancestry State | Lossless | Yes | 32,767 |

### 3.3 Marker alignment

To assess the effectiveness of the Recomb-Mix Cloud pipeline, we applied it to the AoU phased haplotype data for Chromosome 21. We used an eight-way compact reference panel (derived from HGDP and 1000GP) and a recombination map in the GRCh38 build. 200 individuals were randomly selected, and PLINK 2 was used to generate a curated VCF file containing the sites shared between the AoU genomic data and the compact reference panel. Local ancestry inference was then performed using Recomb-Mix with its default parameters. For comparison, we also ran the popular LAI methods FLARE [7] and RFMix [22] following the same procedures.

During the pre-processing step, PLINK 2 curated the genomic dataset by retaining 849,229 biallelic sites from 8,793,215 sites in the original AoU Chromosome 21 phased dataset, overlapping with the 1,891,054 sites in the reference panel. This process was performed in the AoU cloud environment of 32 CPUs and 208 GB RAM, requiring 17,910 seconds of wall-clock time and costing $9.50. During the same process, 200 random individuals were sampled for LAI runs. The ratio of sites in the curated Chromosome 21 dataset to those in the original Chromosome 21 dataset was later used to estimate the number of overlapping sites in other curated chromosomes (see Supplementary Table S5).

To evaluate whether the curated dataset maintains a similar variant distribution to the original, we compared the variant densities of both the original and curated AoU Chromosome 21 datasets. Figure 4 presents a sliding window variant densities plot, where each window displays the proportion of variants, calculated as the number of variants in the window divided by the total variant count across the chromosome. The strong similarity between high- and low-density regions in both datasets demonstrates that the curated dataset retains the variant distribution of the original data. This preservation ensures that the local ancestry inference results remain robust and accurately reflect the ancestry information of the original dataset.

**Figure 4:**
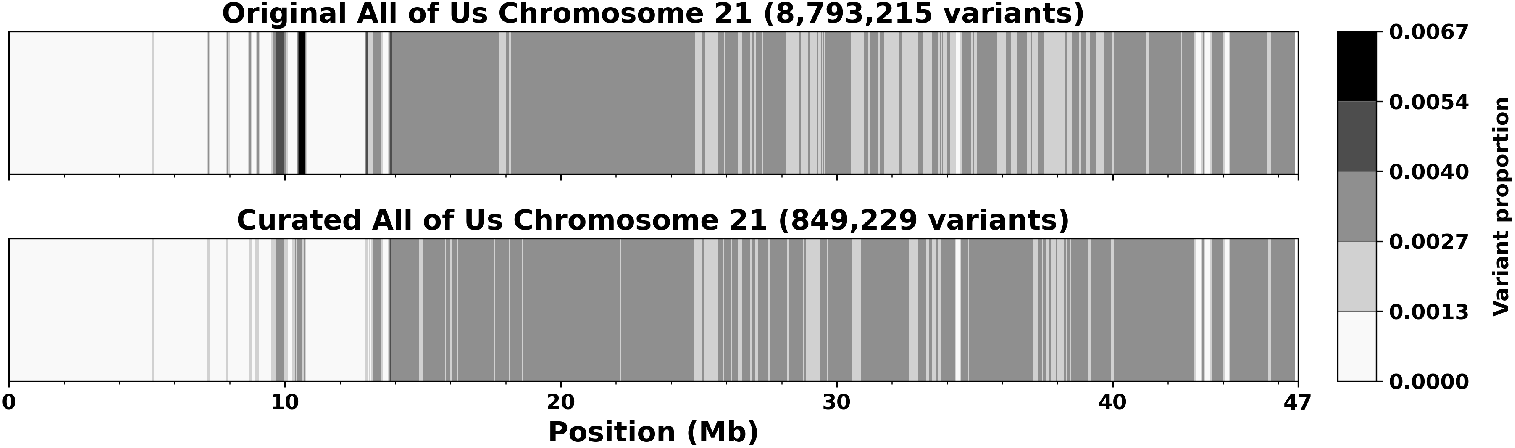
Sliding window variant densities of the original and the curated AoU Chromosome 21 datasets. Window size is 100 kilobase pairs, and step size is 10 kilobase pairs. They are normalized using the variant proportion per window rather than variant counts for comparison. Variant proportion is calculated by dividing the count of variants in a specific window by the total count of variants across the entire chromosome.

### 3.4 Cost

We compared the runtime and the cost of performing LAI for 200 individual samples in the curated AoU Chromosome 21 dataset using Recomb-Mix, FLARE, and RFMix (See Table 3). Subsequently, we estimated the overall cost of scaling pre-processing and LAI runs across all AoU chromosomes for one individual and for all AoU cohorts (See the second-to-last and last columns in Table 3). For Chromosome 21 LAI runs, the same 32 CPUs and 208 GB of RAM environment settings (at a cost of $1.91 per hour) were used. This is because RFMix requires substantial memory, and testing indicated a minimum of 200 GB RAM was needed. To ensure a fair comparison, all programs were run in the same cloud environment setting. All methods utilized parallel processing across the available CPU cores in the cloud environments.

**Table 3:** LAI runtime and cost on 200 individuals from the curated AoU Chromosome 21 dataset, and estimated overall cost (in *Italics*) of performing LAI across all chromosomes for one individual and the entire AoU cohort.

| Method | LAI runtime<br>of 200 Inds for<br>Chr21 (s) | LAI cost of<br>200 Inds for<br>Chr21 (\$) | Overall cost of one<br>Ind for whole<br>genome (\$) | Overall cost of AoU<br>Cohort for whole<br>genome (\$) |
| --- | --- | --- | --- | --- |
| Recomb-Mix | 69 | 0.04 | *0.01* | *4,859.43* |
| FLARE | 2,175 | 1.15 | *0.43* | *179,088.03* |
| RFMix | 23,067 | 12.24 | *4.58* | *1,900,632.53* |

As a result, Recomb-Mix completed its run in just 69 seconds, while FLARE took 2,175 seconds and RFMix took 23,067 seconds. This indicates that for LAI on 200 individuals Chromosome 21 AoU data, Recomb-Mix costs only $0.04, which is about 3% of FLARE’s cost and 0.3% of RFMix’s cost. The overall cost of using the Recomb-Mix Cloud pipeline for 200 individuals for Chromosome 21 data is just $9.54 ($0.05 per individual on average). Cost estimations across all chromosomes and the cohort further emphasize the differences. The estimated pre-processing cost for the whole genome was extrapolated based on the per-variant cost, calculated as $9.50 *×* (658, 219, 451*/*8, 793, 215) = $711.13. Similarly, the estimated LAI cost for all AoU chromosomes was extrapolated based on the per-variant cost, and the estimated LAI cost for all AoU individuals was extrapolated based on the per-individual cost. For example, for Recomb-Mix, the estimated average cost of one individual for the whole genome is calculated as $1.91 *×* (69*/*3, 600) *×* (63, 569, 360*/*849, 229)*/*200 = $0.01, and the estimated cost of the AoU cohort for the whole genome is calculated as $0.01*×* 414, 830 = $4, 148.30. Thus, the estimated overall costs to obtain local ancestry calls for the entire AoU cohort for the whole genome using Recomb-Mix, FLARE, and RFMix are $4, 859.43 (or average $0.01 per individual), $179, 088.03 (or average $0.43 per individual), and $1, 900, 632.53 (or average $4.58 per individual), respectively. Both runtime and cost results demonstrate that Recomb-Mix is the most efficient and cost-effective LAI method and is well-suited for cloud computing environments.

Compared with the runtime cost, the disk storage cost is minimal. For example, AoURW provides 200 gigabytes of persistent cloud storage for $8 per month, which is sufficient to run LAI for 200 individuals on Chromosome 21 using either Recomb-Mix Cloud or the conventional pipeline. However, for studies involving a larger number of individuals or whole-genome analyses, pipeline selection becomes critical. The conventional pipeline can substantially increase storage cost due to the large sizes of the original reference panel and the output file format, which is not optimized for space efficiency.

### 3.5 Ancestry tract length distribution

We evaluated the distributions of ancestry tract lengths generated by three LAI methods applied to the AoU Chromosome 21 dataset. An ancestry tract is a region on a chromosome assigned an ancestry label, and its adjacent regions have different ancestry labels. To do this, we randomly selected 200 individuals from the phased AoU Chromosome 21 data and pre-processed them using the Recomb-Mix Cloud pipeline’s harmonization process. 849,229 markers shared with the eight-way reference panel were identified. We then applied Recomb-Mix, FLARE, and RFMix to this curated dataset to obtain local ancestry calls. All ancestry tracts were collected, and tract length histograms were constructed for each method. For Recomb-Mix and RFMix (version 2), we used their default parameters. For FLARE, we used version 0.6 and set its parameters “min-mac=0” and “min-maf=0” to have the program report as many markers as possible, since its default parameter settings do not report ancestry labels for all markers in the given dataset. It reported 95.91% (814,468 out of 849,229) markers. For markers without reported ancestry labels, we estimated the labels by extending the ancestry label of their nearest neighbor to fill the gap. If both sides reported the same ancestry label, the markers were assigned that label; otherwise, the left neighbor’s ancestry label was assigned.

Figure 5 shows the tract length distributions for the three methods. All three distributions generally follow an exponential distribution pattern, consistent with theoretical expectations [14]. The notable exception is the rightmost bars, which represent unusually large tracts, likely reflecting individuals whose parents are within the same population. We also calculated the percentage of overlapping areas in the histograms for the methods: Recomb-Mix vs FLARE (0.8784), Recomb-Mix vs RFMix (0.9205), and FLARE vs RFMix (0.8957). These results indicate that the tract length distributions are consistent across methods, with Recomb-Mix being more similar to RFMix than to FLARE.

**Figure 5:**
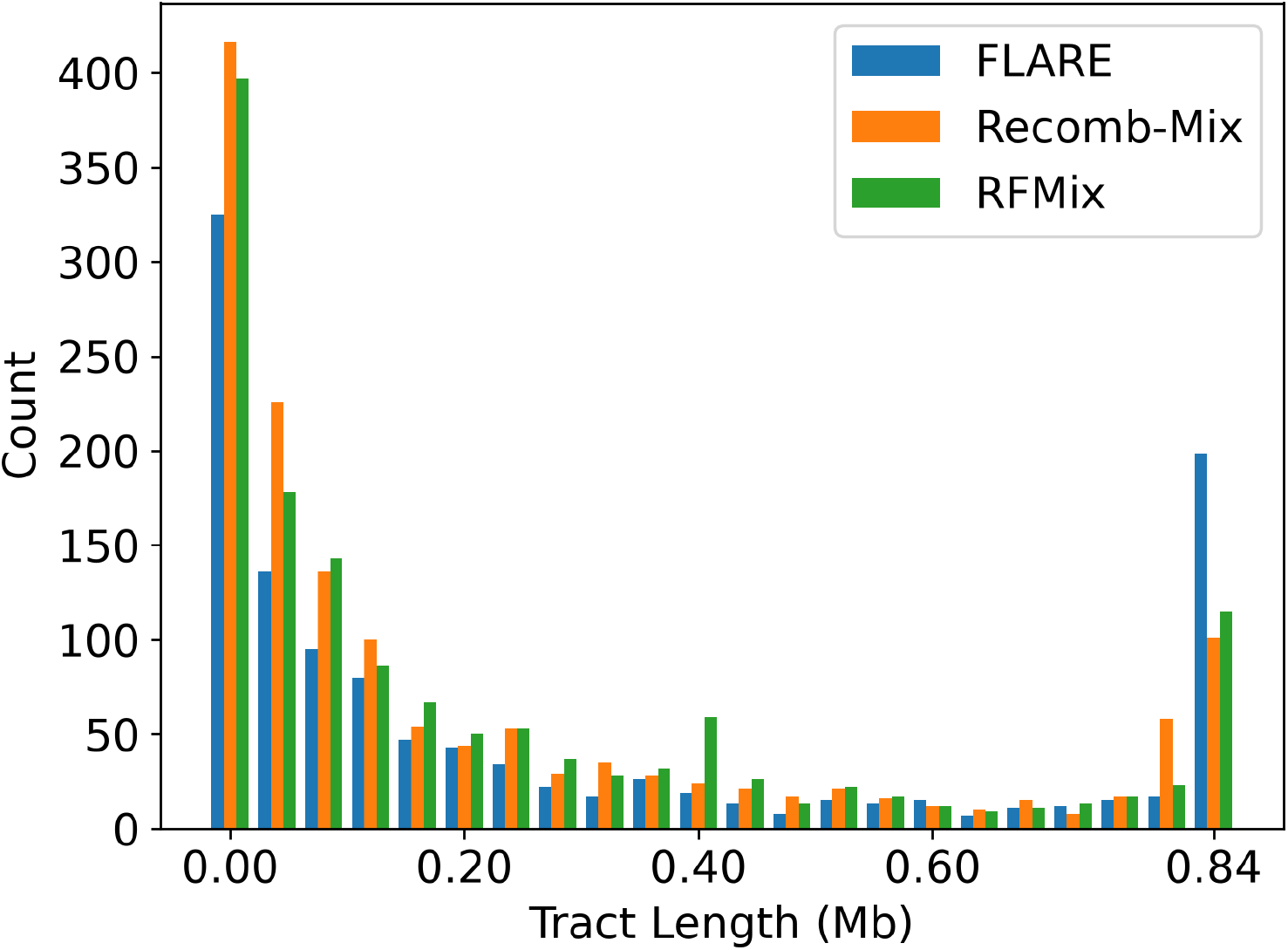
Comparison of the FLARE, Recomb-Mix, and RFMix for ancestry tract length distributions across 200 randomly selected individuals from the AoU Chromosome 21 dataset. The bin size is 40,000 base pairs. An eight-way reference panel is used for all three LAI methods, and 849,229 markers are found shared with the AoU data. Recomb-Mix and RFMix were run with their default parameters. FLARE was configured with parameters min-mac=0 and min-maf=0 to ensure its results covered as many regions as possible. For regions without ancestry calls from FLARE (4.09%), ancestry calls were estimated by copying those from adjacent regions.

## 4 Discussion

We presented Recomb-Mix Cloud, a cost-efficient and cloud-friendly pipeline designed to streamline local ancestry inference for diverse research applications. By deploying and running the pipeline on the AoU platform as an example, we demonstrated its effectiveness at delivering reliable results with minimal computational cost in the cloud environment. Our pipeline removes the burden from end users by incorporating complex steps of data preprocessing and LAI performing, making the process accessible to those with limited expertise in the area.

Our pipeline leverages compact reference panels, which significantly reduce storage requirements compared to traditional panels while preserving robust ancestral signals. The compact reference panel enables the power of populations with small sample sizes to be represented as references. For end users, we provide pre-computed compact reference panels across a variety of population combinations and reference builds, as well as support the creation of customized reference panels tailored to their research needs.

To further enhance usability, we also provide a LAI output format conversion utility to easily convert inferred local ancestry calls into multiple output formats. This flexibility facilitates downstream analyses by allowing end users to integrate results easily into their workflows and pipelines, accelerating research progress and fostering broader adoption of local ancestry inference in research studies.

While the Recomb-Mix Cloud pipeline is cost-efficient, there are still challenges to address. One challenge is the insufficient representation of ancestries in the standard reference panels. These panels were composed of data from the HGDP and 1000G datasets, which provide publicly available genomic data from diverse populations across multiple continents. However, they do not fully capture the human genetic diversity. Many small, isolated, or intra-continental-level populations remain underrepresented in these datasets, and are actively being sequenced and studied (e.g., indigenous Americans [26]). Since they were absent from the standard reference panel, the LAI process would normally assign nearby ancestry-group labels to them based on geographic proximity, which may not yield the most accurate results.

Another challenge is that our pipeline’s performance depends on the percentage of variants in the original reference panels that overlap with those in the target dataset. The available biobank-scale sequencing dataset in the cloud typically includes variants not present in the reference panels. For example, the AoU Chromosome 21 dataset has only 9.66% of variants overlapping with those in the reference panel, which is constructed from the HGDP and 1000G datasets. Thus, the LAI result from the pipeline reflects a merely small portion of the target genome. Though ancestry labels for variants not present in the reference panel can still be estimated, they are rather preliminary and may affect downstream analysis outcomes.

## Data availability

The Recomb-Mix Cloud pipeline and the compact reference panels are freely available at https://github.com/ucfcbb/Recomb-Mix.

## Acknowledgments

This work was supported by the National Institutes of Health under grants R01HG010086 and R01AG081398.

