## Supplementary Materials for "Recomb-Mix Cloud: a cost-efficient and scalable cloud-friendly pipeline for local ancestry inference"

| Section | Element | Type | Description |
| --- | --- | --- | --- |
| Result | physical_position | integer | Base coordinate of a chromosome |
| Result | individual_haplotype_ancestry_identifier | integer | Ancestry indicator of an individual haplotype for the physical_position |

Table S1: The ancestry dosage format schema.

| Section | Element | Type | Description |
| --- | --- | --- | --- |
| Result | physical_position | integer | Base coordinate of a chromosome |
| Result | individual_diploid_ancestry_dosage | integer | Ancestry count of an individual for the physical_position |

Table S2: The ancestry state format schema.

| Scenarios | Recomb-Mix | FLARE | RFMix |
| --- | --- | --- | --- |
| Gen-15 Ref-100 | 0.9440 | 0.1789 | 0.3796 |
| Gen-15 Ref-250 | 0.9564 | 0.4096 | 0.7975 |
| Gen-15 Ref-500 | 0.9697 | 0.6603 | 0.8953 |
| Gen-15 Ref-1000 | 0.9731 | 0.8017 | 0.9423 |
| Gen-50 Ref-500 | 0.8611 | 0.3232 | 0.5404 |
| Gen-100 Ref-500 | 0.7746 | 0.2020 | 0.2929 |
| Gen-200 Ref-500 | 0.5216 | 0.0855 | 0.1315 |

Table S3: The mean diploid squared Pearson's correlation coefficient $r^2$ of three-way inter-continental simulated Chromosome 18 datasets on Recomb-Mix, FLARE, and RFMix under seven scenarios: 15-generation dataset with the reference panel sizes 100, 250, 500, and 1,000, and 50-, 100-, and 200-generation datasets with the reference panel size 500.

| Chromosome ID | Number of Markers | Original File Size | Compact File Size |
|---|---|---|---|
| 1 | 10,849,042 | 1,488.17 | 92.48 |
| 2 | 11,201,075 | 1,498.57 | 95.39 |
| 3 | 9,136,229 | 1,256.48 | 78.13 |
| 4 | 8,946,351 | 1,254.21 | 76.23 |
| 5 | 8,316,361 | 1,133.33 | 70.99 |
| 6 | 7,672,616 | 1,110.84 | 65.71 |
| 7 | 7,514,152 | 1,058.48 | 64.26 |
| 8 | 7,117,448 | 964.61 | 60.69 |
| 9 | 5,993,088 | 843.17 | 51.08 |
| 10 | 6,363,090 | 910.34 | 54.48 |
| 11 | 6,444,957 | 889.41 | 54.87 |
| 12 | 6,150,120 | 854.83 | 52.64 |
| 13 | 4,829,705 | 687.74 | 41.18 |
| 14 | 4,118,981 | 571.83 | 35.37 |
| 15 | 3,857,373 | 540.06 | 33.12 |
| 16 | 4,329,890 | 579.88 | 37.02 |
| 17 | 3,770,988 | 519.19 | 32.46 |
| 18 | 3,742,180 | 519.35 | 31.91 |
| 19 | 2,886,968 | 410.94 | 25.06 |
| 20 | 3,093,286 | 456.03 | 26.48 |
| 21 | 1,891,054 | 291.35 | 16.10 |
| 22 | 1,982,340 | 304.98 | 17.12 |

Table S4: The number of markers and sizes (megabytes) of the eight-way inter-continental original and compact reference panels in GRCh38 build.

| Chromosome ID | Number of Markers in Original Dataset | Number of Markers in Curated Dataset |
|---|---|---|
| 1 | 52,886,684 | *5,107,677* |
| 2 | 58,252,492 | *5,625,895* |
| 3 | 48,174,243 | *4,652,560* |
| 4 | 46,539,832 | *4,494,713* |
| 5 | 43,368,264 | *4,188,410* |
| 6 | 40,692,546 | *3,929,995* |
| 7 | 38,094,802 | *3,679,111* |
| 8 | 36,883,753 | *3,562,150* |
| 9 | 29,280,982 | *2,827,892* |
| 10 | 32,006,801 | *3,091,145* |
| 11 | 32,555,692 | *3,144,156* |
| 12 | 31,406,215 | *3,033,142* |
| 13 | 23,301,209 | *2,250,379* |
| 14 | 21,290,034 | *2,056,144* |
| 15 | 19,430,474 | *1,876,552* |
| 16 | 21,104,532 | *2,038,228* |
| 17 | 18,600,120 | *1,796,358* |
| 18 | 18,230,491 | *1,760,660* |
| 19 | 13,873,668 | *1,339,888* |
| 20 | 14,621,774 | *1,412,138* |
| 21 | 8,793,215 | 849,229 |
| 22 | 8,831,628 | *852,939* |

Table S5: Number of markers in the original and curated AoU datasets. The estimated numbers (in *Italics*) were estimated from the ratio of markers in the curated Chromosome 21 dataset to those in the original Chromosome 21 dataset. The number of markers in the original AoU data is 658,219,451, and the estimated number of markers in the curated AoU data is *63,569,360*, respectively.
